# Psilocybin acutely reduces low-frequency BOLD power and frequency-specific connectivity

**DOI:** 10.64898/2026.04.09.717379

**Authors:** Anders S. Olsen, Kristian Larsen, Drummond E-W. McCulloch, Melanie Ganz, Martin K. Madsen, Brice Ozenne, Gitte M. Knudsen, Naveed ur Rehman, Patrick M. Fisher

## Abstract

Psilocybin and other serotonergic drugs acutely alter human brain function and large-scale connectivity as measured with BOLD fMRI, but whether these effects are frequency-specific remains unknown. We applied multitaper spectral and cross-spectral analyses to resting-state fMRI data from 28 healthy volunteers scanned multiple times acutely following oral psilocybin administration (0.2 – 0.3 mg/kg), together with plasma psilocin measurements, to estimate psilocin associations with temporal frequency-specific activity and connectivity. Psilocybin produced a selective reduction in low-frequency spectral power (0.01 – 0.06*Hz*) and an increase in spectral entropy, with the strongest effects in transmodal networks. We also observed a reduction in low-frequency connectivity energy explained by the unimodal/transmodal axis. These findings demonstrate that psilocin induces spatially distributed, frequency-dependent alterations, suggesting that broadband fMRI analyses may obscure low-frequency dynamics. Frequency-resolved approaches may offer greater sensitivity for characterizing psychedelic effects on brain activity.

## 1 Introduction

Psychedelic compounds such as psilocybin elicit profound changes in perception, cognition, and self-experience (Carhart-Harris et al., 2012; Madsen et al., 2020; McCulloch et al., 2022; Roseman et al., 2017). Human electro- and magnetoencephalography (EEG/MEG) studies show reductions in broadband spectral power following psychedelic administration (Alamia et al., 2020; Ip et al., 2026; Murray et al., 2022; Muthukumaraswamy et al., 2013; Schenberg et al., 2015; Timmermann et al., 2023), with particularly strong effects in the alpha band (Godfrey et al., 2025). Complementary fMRI work has reported changes in signal complexity (McCulloch et al., 2024) and large-scale network organization, most consistently finding reduced canonical network integrity together with increased global integration (Madsen et al., 2021; Tagliazucchi et al., 2016). However, the frequency-specific structure of BOLD activity and interregional coupling under psychedelics has received far less attention.

BOLD signals are dominated by structured, spontaneous fluctuations at low frequencies (commonly ∼ 0.01 − 0.08*Hz*), yet different brain regions and networks can preferentially express power and coupling at distinct spectral bands (Wu et al., 2008; Zang et al., 2007). Several studies suggest psychedelics reduce broadband spectral power of low-frequency hemodynamic signals after psychedelics in both animals and humans (Barrett et al., 2020; Delli Pizzi et al., 2023; Faramarzi et al., 2024; Golden and Chadderton, 2022; La-Torraca-Vittori et al., 2025; Subramanian et al., 2025; Tagliazucchi et al., 2014; Wall et al., 2023). However, all these studies evaluated the amplitude of low-frequency fluctuations (ALFF), a broadband summary statistic that does not consider frequency-specific power changes following psychedelic administration. As such, they implicitly assume that psilocybin effects are spread evenly over a broad frequency band, most commonly 0.01 − 0.08*Hz*. Instead, determining psilocybin effects on the spatiospectral power distribution and the frequency-resolved patterns of functional coupling can provide a richer and more nuanced account of altered brain states *vis-`a-vis* conventional broadband power and connectivity metrics.

A persistent practical challenge for psychedelic fMRI is head motion, which produces nonlinear, broad-band artifacts that can mimic the hallmarks of altered connectivity (for example, apparent increases in global connectivity) and often survive standard realignment and linear denoising pipelines (Ciric et al., 2017; Power et al., 2012, 2017; Satterthwaite et al., 2013; Van Dijk et al., 2012). This confound is especially problematic for spectral analysis because motion artifacts span many frequencies depending on their temporal morphology and recurrence pattern. To address these issues, we adopt a frequency-specific connectivity analysis that emphasizes narrowband structure while controlling for broadband, noise-dominated components. Concretely, we apply the generalized eigendecomposition (GED, Parra and Sajda, 2003) to cross-spectral density matrices to isolate connectivity patterns that maximally differentiate narrow-band signal from broadband background, which provides resilience to motion-related contamination and represents a principled way to extract frequency-specific networks (Cohen, 2022; Rosso et al., 2025).

One of the effects of psilocybin, supported by multiple studies, is increased brain entropy, i.e., a measure of randomness in the brain (Carhart-Harris, 2018; Carhart-Harris et al., 2014; McCulloch et al., 2024). Our lab recently evaluated 13 brain entropy metrics on the dataset investigated herein, finding diverse results, indicating that “entropy” is not a single construct and should instead be treated as a summary statistic affected by several underlying brain functions (McCulloch et al., 2024). Here we estimate *spectral entropy*, a measure of the flatness of the BOLD power spectrum, to complement spatiospectral analyses with a description of psilocybin effects along a dimension of information.

We apply these methods to a densely sampled resting-state fMRI dataset wherein *N* = 28 healthy participants completed scans at baseline and multiple time points during the acute psychedelic experience following oral psilocybin administration, coupled with plasma psilocin level (PPL) data (Madsen et al., 2021; McCulloch et al., 2024). First, we map regional spectral power across BOLD frequencies to quantify how psilocybin alters the spatiospectral landscape of BOLD fluctuations. Second, we apply GED to derive frequency-specific connectivity patterns that are maximally different from broadband noise and motion effects. Together, these approaches aim to (i) identify fingerprints of the psilocybin state in (univariate) regional BOLD spectral power and (ii) reveal which frequency-resolved connectivity patterns are selectively modulated by psilocybin. Combined, we demonstrate an analytic framework for studying pharmacological perturbations of frequency-specific BOLD dynamics, thereby advancing our understanding of how serotonergic psychedelics reorganize large-scale brain function.

## 2 Methods

### 2.1 Study description

We re-analyzed data from a larger study on human serotonin 2A receptor modulation (NCT03289949) previously described in detail elsewhere (Madsen et al., 2021; McCulloch et al., 2024) with further analyses in (Larsen et al., 2025; Olsen et al., 2022). The study was approved by the ethics committee for the capital regions of Denmark (journal identifier: H-16028698) and Danish Medicines Agency (eudraCT identifier: 2016-004000-61), and all participants provided written informed consent. In short, *N*_*subs*_ = 28 participants received either peroral psilocybin (0.2-0.3 mg/kg) or ketanserin (data not included here) in a single-blind cross-over study. Resting-state fMRI data were acquired on two Siemens 3T Prisma scanners (scanner 1: TR=0.8s (multiband, 8x acceleration, N=13), scanner 2: TR=2s (N=15)) pre-drug and multiple times during the acute phase up to 350 minutes after drug administration (Fig. 1A), 1-10 scans per subject. All fMRI-scans were 10 minutes long except five subjects on scanner 2, who were scanned with 5-minute sequences in two different phase-encoding directions. For further details regarding imaging sequences, see McCulloch et al., 2024. Of 170 total scans, four were discarded for being interrupted, and one was discarded for having the wrong coils active.

**Figure 1.**
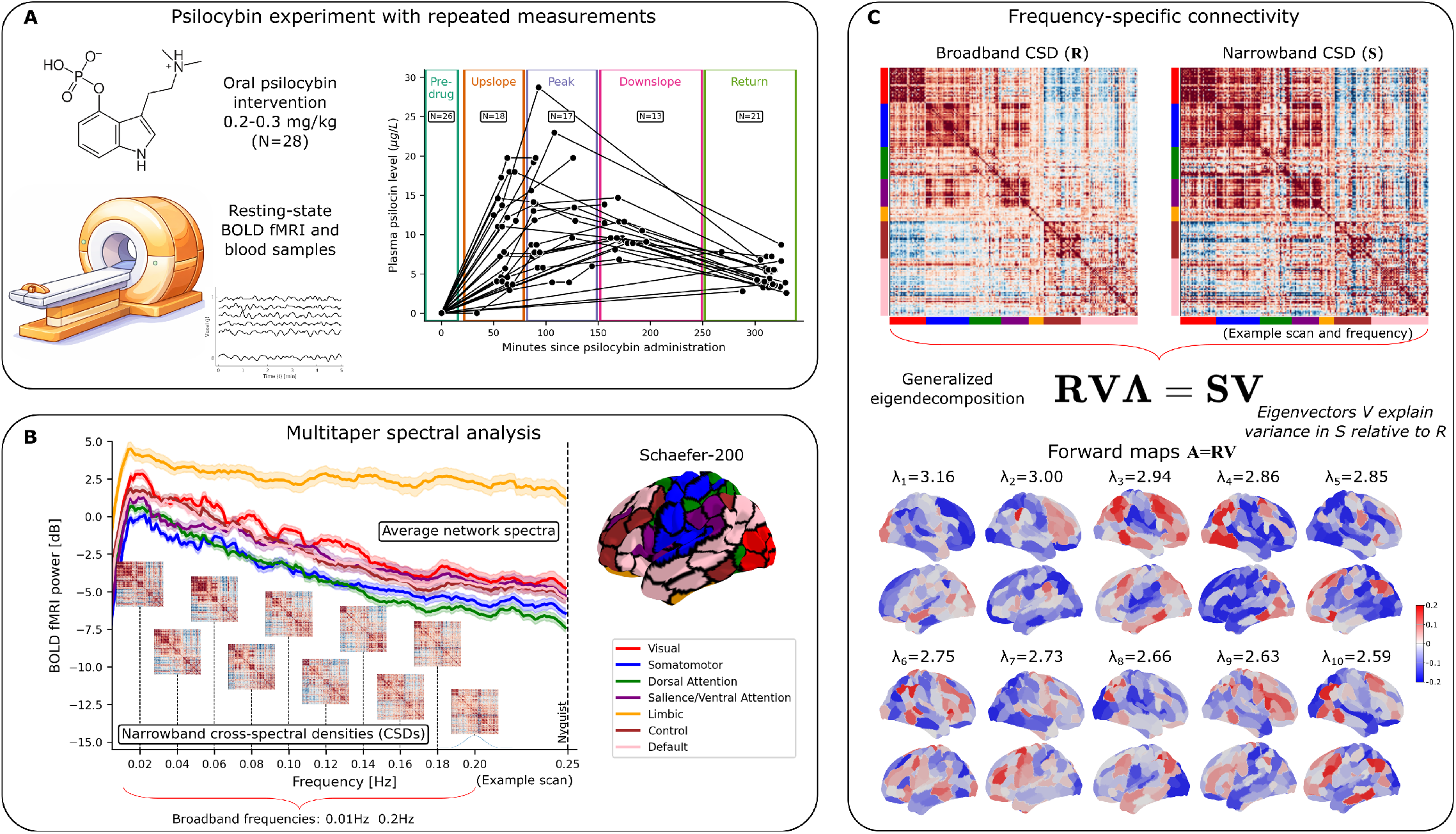
Methods overview for frequency-specific assessment of BOLD fMRI activity and connectivity. **A**): N=28 participants were scanned in resting-state several times during the acute phase of a psilocybin experiment with concurrent blood samples for the plasma psilocin level. Only included scans following quality control are visualized. **B**) Each voxel time series was converted to a power spectrum using multitaper spectral analysis and subsequently parcellated. Narrowband cross-spectral density (CSD) matrices were computed on parcellated time series using multitaper cross-spectral estimation and averaged using a Gaussian frequency filter. The broadband CSD was computed in the frequency window from 0.01 − 0.20*Hz*. “Nyquist” indicates the maximum frequency at which spectral power can be estimated. **C**) The connectivity patterns maximally explaining the contrast between narrowband and broadband CSD were extracted using the generalized eigendecomposition (GED). While the GED eigenvectors **V** are not directly phsiologically interpretable, the forward maps **A** = **RV** were evaluated for each scan and frequency. Only the leading 10 forward maps were retained.

Each scan was matched with a blood measurement of psilocin (PPL), i.e., the psychoactive substance, which is directly related to both the intensity of the subjective experience and cortical serotonin 2A receptor occupancy (Madsen et al., 2021). Blood sample timings were matched with the midpoint of the fMRI-scan; missing PPL values before drug administration were imputed to zero (N=1). If the scan was acquired *<* 200 minutes after drug administration, it was included if a blood sample was acquired no more than 20 minutes before/after the fMRI scan. If the scan was acquired after 200 minutes, it was included if a blood sample was acquired no more than 40 minutes before/after the fMRI scan. Based on these criteria, one scan was excluded.

We used “fMRIprep” (v25.1.3, defaults, Markiewicz et al., 2026) for fMRI preprocessing into fsLR-32k standard space, containing a surface representation for the cortex. Through quality control, we discarded an additional 12 scans across five subjects for poor image registration and corrupted fieldmaps. We quantified motion summary statistics for each scan: mean framewise displacement (FD), max FD, and pct-high-motion (ratio of volumes with FD*>*0.5mm). Motion-heavy scans (maxFD*>*3mm or pct-high-motion*>*20%) were discarded, leaving a final 123 scans for analysis. 6 motion parameters and a mean white matter, gray matter and cerebrospinal fluid time series (totaling nine components) were regressed out using “nilearn” (v0.13.1, contributors, n.d.) to reduce residual non-neural physiological signal. Spectral power is sensitive to the scale of the time series. However, z-scoring time series would imply that we could only infer relative shifts within frequency spectra due to Parseval’s theorem. Instead, we rescaled time series to “percent change” to reduce spatial differences in BOLD mean and variance without normalizing them. To reduce the effect of noise in subsequent spatial downsampling using parcellations, vertices with low temporal variation (tSNR *<* 30), large artifacts (any volume with percent change *>* 25%), or non-Gaussian distribution (Fisher’s kurtosis *>* 10) were excluded.

For visualization purposes, the 123 scans were sometimes subdivided into five phases of the acute psilocybin experience depending on their time of acquisition relative to drug administration (Fig. 1A): “Predrug” (*N* = 26, *N*_*scans*_ = 33), “Upslope” (0-80 minutes, *N* = 18, *N*_*scans*_ = 23), “Peak” (80-150 minutes, *N* = 17, *N*_*scans*_ = 25), “Downslope” (150-250 minutes, *N* = 13, *N*_*scans*_ = 17), “Return” (after 250 minutes, *N* = 21, *N*_*scans*_ = 27). Notably, this subdivision was not used for statistical analysis, only to assist relevant visualizations.

### 2.2 Multitaper spectra

To estimate univariate spectra and pairwise cross-spectra we used multitaper spectral estimation (Thomson, 1982) as implemented in “nitime” (v0.11, Rokem et al., 2009) (Fig. 1B). Multitaper methods improve spectral bias/variance properties over alternatives, e.g., Welch and Bartlett, and are particularly beneficial for timeseries with short acquisition lengths relative to the longest wavelength (Mitra and Pesaran, 1999). The multitaper spectral estimate is

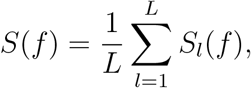

where *S*_*l*_(*f*) is the periodogram (squared absolute Fourier transform) of the input data *x*(*t*) multiplied element-wise by a Slepian function *g*_*l*_(*t*). We used normalized half-bandwidth *NW* = 3, corresponding to *L* = 5 Slepian tapers. Some scans were *T* = 300 seconds long, meaning that the lowest frequency we can reliably quantify using Slepian tapers is *f*_*min*_ = *NW/T* = 0.01*Hz*. Univariate spectra were estimated for each cortical vertex and subsequently parcellated into *p* = 200 cortical parcels (Schaefer et al., 2018) grouped into 7 cortical networks (Yeo et al., 2011). All nitime function calls used default parameters except for NFFT, which was 2.5 times larger for the short-TR scanner to account for their difference in TR.

We performed two statistical analyses of the relation between spectral power and PPL. The first analysis was performed on seven cortical networks (Yeo et al., 2011) in full frequency resolution (0.01*Hz* spacing). The second analysis was performed on 200 cortical parcels in three canonical frequency bands: Slow-3 (0.067 − 0.2*Hz*), Slow-4 (0.025 − 0.067*Hz*), and Slow-5 (0.01 − 0.025*Hz*, Buzsáki and Draguhn, 2004), thus increasing the spatial resolution while reducing the frequency resolution.

#### 2.2.1 Statistical analysis of spectral power

For each network and frequency spaced by 0.01*Hz*, we fitted spectral power in dB, i.e., 10 log_10_(*S*(*f*)), using a linear mixed effects model with PPL, age, sex, scanner, and number of volumes as fixed effects and subject as random intercept. We also included motion-related nuisance regressors: mean FD, max FD, and pct-high-motion, representing three complementary characterizations of motion. The p-value for PPL was calculated as a likelihood ratio (LR) test between a model with PPL and a model without PPL. Reported partial correlation coefficients were calculated as the Pearson correlation coefficient between PPL and the residuals of the statistical model estimate without PPL.

To control the family-wise error rate and account for multiple testing across 20 frequency bins between 0.01*Hz* and 0.20*Hz*, we performed permutation testing with max-T adjustment by scrambling the normalized residuals of each linear mixed effects model 1000 times (Lee and Braun, 2012). Briefly, for each permutation, all 20 models were re-estimated using the scrambled residuals, and the maximum likelihood ratio was retained. Repeating for all permutations, this led to a pool of 1000 “max-LR” values. The p-value (*p*_max-T_) for the relation between PPL and dB was calculated for each frequency as the number of instances the permuted max-LR values were greater than the observed LR, divided by 1000. Subsequently, we performed Bonferroni-correction across 7 networks, i.e., *p*_FWEℝ_ = 7 *× p*_max-T_.

In a second analysis, with higher spatial resolution but only three frequency bands, the linear model formulation was equivalent, but p-values were corrected using max-T across parcels and Bonferroni-corrected across frequency bands. All statistical testing was performed with the R-software (R Core Team, 2023) using the “nlme” package (Pinheiro and Bates, 2000).

#### 2.2.2 Spectral entropy

Using the univariate spectra, we also quantified spectral entropy as the Shannon entropy

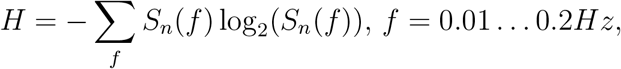

Where 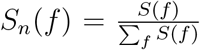. The entropy, measured in *bits*, is a metric of the spread of states in a system and spectral entropy is therefore a measure of spectral *flatness*. Spectral entropy is maximized for white noise and minimized (zero) for a single-frequency sine wave. Vertex-wise entropy values were parcellated and p-values were estimated using max-T correction across parcels.

### 2.3 Generalized eigendecomposition

The GED (Parra and Sajda, 2003) is an analysis tool that can be used to maximally explain axes of variance in one covariance matrix relative to another. The GED has previously been used for EEG/MEG data to maximally separate temporal task segments from rest in a block-design experiment (Cohen, 2022) and frequency-specific spatial covariance from broadband, 1/f background signal (Rosso et al., 2025). Here, we apply GED to cross-spectral densities (CSDs, the frequency-domain equivalent of covariance) and frequency-resolved fMRI data, neither of which have been explored in the literature, to the best of our knowledge. Evaluating the spatial covariance in the frequency domain rather than temporally filtered covariance in the time domain allows for directly controlling the bandwidth (filter response) at each frequency and eliminates a computationally heavy processing step. We assume that head motion, which is particularly problematic in psychedelic brain imaging studies, is captured entirely by the broadband CSD matrix. As such, the narrowband generalized eigenvectors will be motion-free, since motion is directly controlled for via the normalization by the broadband CSD. As such, the GED is ideal for evaluating frequency-specific connectivity effects of psilocybin while mitigating confounding motion effects.

We first computed the multitaper CSDs for each pair of cortical regions using parcellated time series as input (Fig. 1C). The CSDs were estimated using “nitime” (v0.11, Rokem et al., 2009) with the same parameters as the univariate power spectra and then downsampled to 0.01*Hz* spacing using a Gaussian filter with variance *σ*^2^ = 0.01*Hz* to obtain a set of narrowband CSDs **S**_*f*_ ∈ ℝ^*p×p*^, *f* = 0.01, …, 0.2*Hz*. We also computed the average CSD over these frequencies to produce the broadband CSD **R** ∈ ℝ^*p×p*^. Subsequently, we computed the GED (dropping the frequency subscript)

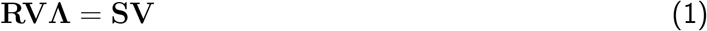

where **V** are eigenvectors that maximally explain variance in the “ratio” between the narrowband and broadband covariance **R**^−1^**S**. When **R** and **S** are of the same scale, eigenvalues in the diagonal of **Λ** are directly interpretable as a ratio of variances (generalized Rayleigh quotients)

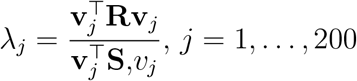

i.e., if *λ*_*j*_ *>* 1, *j* = 1, …, *p* the corresponding eigenvector **v**_*j*_ explains *more* variance in **S** than in **R** (Cohen, 2022). Here we ensured equiscaling by trace-normalizing both **R** and **S** prior to calculating the GED. For numerical stability, **R** was shrinkage-regularized by adding 0.01 to all diagonal elements and downweighting all off-diagonal elements by the same amount.

Eigenvectors in the columns of **V** are orthonormal in *broadband-space*, i.e.,

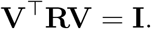

As such, eigenvectors **V** are not directly comparable across scans and subjects. Furthermore, as detailed in Haufe et al., 2014, the eigenvector represents a filter, which is not directly physiologically interpretable. Instead, the eigenvectors should be weighted by the data, i.e., either by the narrowband CSD **SV** or broadband CSD **RV**. Given Eq. (1), **RV** = **SVΛ**^−1^, i.e., these two representations are equivalent, save for column scaling by eigenvalues **Λ**. As such, **SV** is comparable to a covariance map scaled by the units of the input data, whereas **RV**, which corresponds to the forward model representation by (Haufe et al., 2014), represents the topography shape without being dominated by amplitude differences between columns or across scans/subjects. Here we focus on topography maps **A** = **RV** and retain the leading 10 columns, which empirically all explained more narrowband than broadband covariance, i.e., *λ*_*j*_ *>* 1, *j* = 1, …, 10.

#### 2.3.1 Generalized Procrustes analysis

Since eigenvectors are sign-invariant, the spatial maps **A** are also of arbitrary sign. Furthermore, columns are ordered by decreasing eigenvalues, the order of which may change between scans or frequencies. As such, averaging or modeling sets of **A**-matrices arithmetically without ensuring consistent signs and column ordering risks smearing out information. Here we suggest an automatic approach using the generalized Procrustes analysis (GPA) (Dryden and Mardia, 2016; Gower, 1975), which is an iterative averaging method using the Procrustes alignment problem.

Given two matrices **A**_1_, **A**_2_ ∈ ℝ^*p×q*^, the original Procrustes problem concerns finding the rotation matrix **Q** that best maps **A**_2_ to **A**_1_ (Schönemann, 1966)

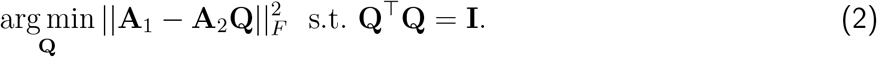

The Procrustes problem is particularly useful in shape analysis (Dryden and Mardia, 2016), wherein objects should be compared regardless of translation, rotation, and reflection. The problem is solved using the thin singular value decomposition (SVD)

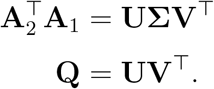

For a set of matrices 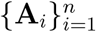, e.g., one per scan or frequency, the GPA starts with an arbitrarily chosen representative sample **A**^(*r*)^ ∈ ℝ^*p×q*^ and then iterates between 1) aligning all samples to the representative using Eq. (2) to obtain 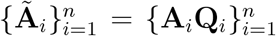, and 2) computing the new average using the aligned matrices. The algorithm ran until an absolute Frobenius-norm difference of 10^−9^ between consecutive iterations. The algorithm is sensitive to the initialization but reasonably fast; here we evaluated the result after initializing with all possible scans and analyzed the solution with the lowest summed Frobenius norm 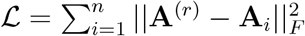.

In our analysis, we first used GPA to align forward maps across scans, producing a set of 10 average maps for each frequency. Secondly, we used GPA to align average maps across frequencies, producing a set of 10 *grand mean* maps. To evaluate how well these grand-mean forward maps mapped back to individual narrowband CSDs, we computed the Rayleigh quotient

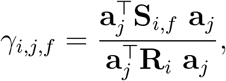

where *i* = 1, …, *n* indexes fMRI scans, *j* = 1, …, 10 indexes grand-mean forward maps, and *f* indexes frequencies. To obtain an indication on how the grand-mean forward maps explained broadband data, we also computed the Rayleigh quotient for broadband CSDs:

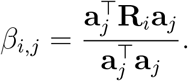

We related Rayleigh quotients, as well as GED eigenvalues corresponding to the leading eigenvector to PPL statistically using the same linear model formulation as previously, including max-T correction across frequencies. Rayleigh quotients were additionally Bonferroni-corrected across 10 average forward maps, i.e., *p*_FWER_ = 10 *× p*_max-T_.

## 3 Results

### 3.1 Univariate power spectral effects of psilocybin

We computed multitaper spectral power of BOLD fMRI vertex-wise time series, summarized spectral power in seven cortical networks (Yeo et al., 2011), and evaluated the relation between log-power and plasma psilocin level (PPL) in a linear mixed effects model. Statistically, we observed widespread significant negative associations between PPL and spectral power at frequencies 0.01−0.06*Hz* (Fig. 2A). These were strongest in the control network (*f* = 0.01*Hz*: partial correlation [CI] *r* = −0.44 [−0.57, −0.29], *p*_FWER_ *<* 0.001; *f* = 0.06*Hz*: *r* = −0.26 [−0.41, −0.08], *p*_FWER_ *<* 0.001) followed by, in order, default mode, dorsal attention, salience, visual, and somatomotor networks. Only the limbic network had no significant associations to PPL.

**Figure 2.**
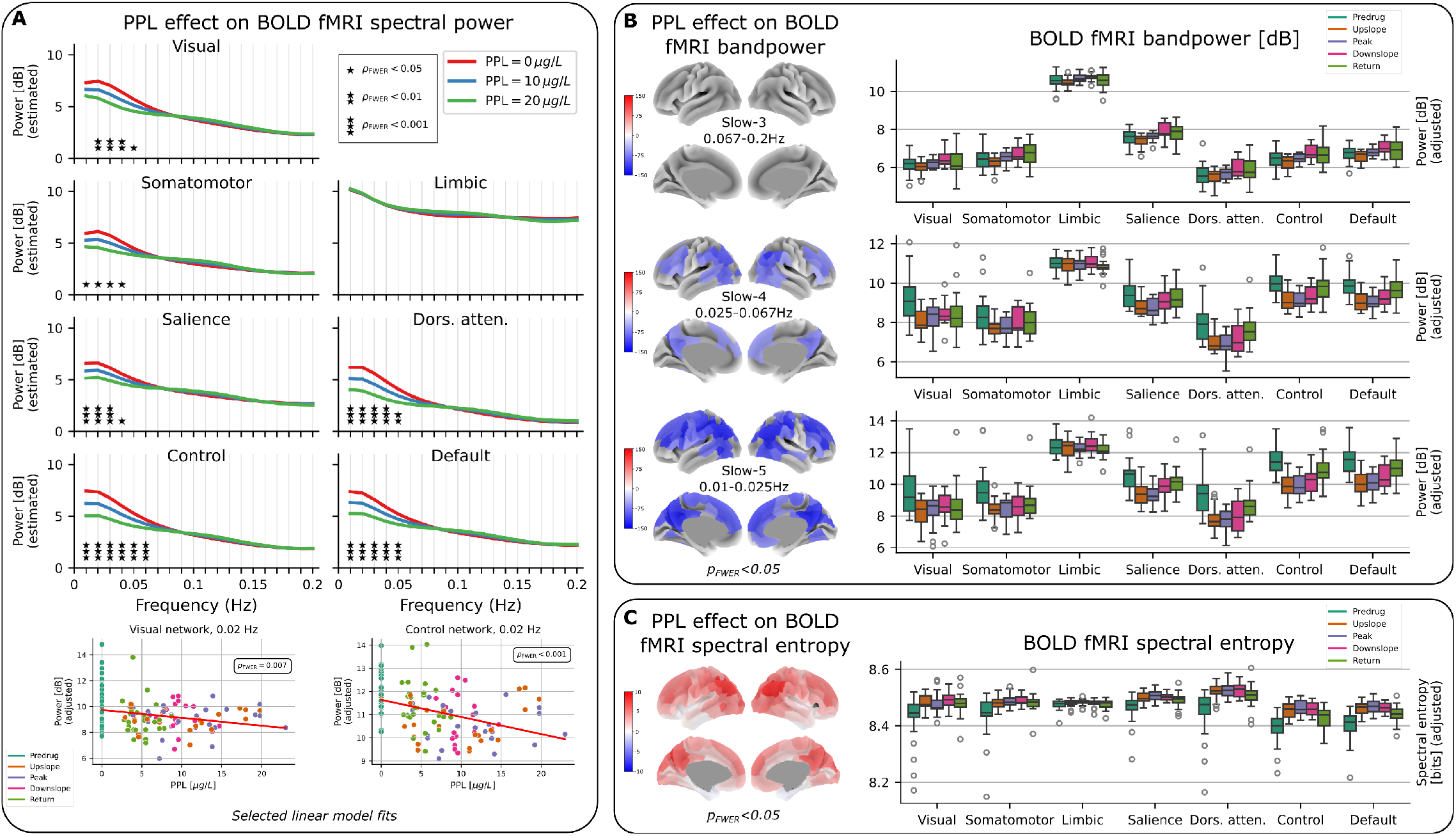
Univariate spectral power analysis of BOLD fMRI data and relation to psilocybin. **A**) Estimated log-power as a function of plasma psilocin level (PPL) from a mixed effects model. Asterisks indicate corrected statistical significance for PPL. **B**) Spectral power analysis in 200 parcels and 3 frequency bands. Brain maps show the linear model coefficient for PPL for parcels with *p*_*F W ER*_ *<* 0.05. Boxplots show the spectral power across subjects within time intervals after regressing out nuisance variables (i.e., “adjusted”); if a subject had more than one scan per time-interval, these were averaged. **C**) Same as B) for spectral entropy.

In a secondary analysis, we increased the spatial resolution to 200 parcels (Schaefer et al., 2018) and reduced the spectral resolution to three frequency bands (Fig. 2B). We observed no significant associations between log-power and PPL in slow-3 (0.067 − 0.2*Hz*). Instead, power in both slow-4 (0.025 − 0.067*Hz*) and slow-5 (0.01 − 0.025*Hz*) was statistically significantly negatively associated with PPL in widespread high-order association cortices, strongest in lateral parietal cortex (left hemisphere, slow-5: *r* = −0.49 [−0.61, −0.34], *p*_FWER_ *<* 0.001), posterior cingulate (left hemisphere, slow-5: *r* = −0.46 [−0.59, −0.31], *p*_FWER_ *<* 0.001) and medial prefrontal cortices (left hemisphere, slow-5: *r* = −0.46 [−0.59, −0.32], *p*_FWER_ *<* 0.001). Boxplots for salience, dorsal attention, control, and default mode networks in these frequency ranges visually indicated a U-shape pattern, wherein the distribution of spectral power across subjects for most networks were highest for predrug, lowest for upslope, peak, and downslope, and close to the predrug levels for the return phase. Spectral power for the return phase was still lower than predrug, indicating a drug effect reaching beyond the ∼ 350 minutes of data acquisition. For visual and somatomotor networks, the return phase did not display visually higher power than upslope/peak/downslope, indicating a more persistent drug effect.

We observed statistically significant positive associations between PPL and spectral entropy in the same association areas as above (left lateral parietal cortex: *r* = 0.49 [0.34, 0.61], *p*_FWER_ *<* 0.001; Fig. 2C). Similarly, boxplots visually displayed an inverted U-shape, indicating stronger effects for transmodal than unimodal networks.

### 3.2 Frequency-specific connectivity effects of psilocybin

We computed generalized eigenvectors, i.e., the spatial filters that maximally explained narrowband vs broadband connectivity in each resting-state fMRI scan, yielding a set of 10 generalized eigenvalues per frequency and scan. Fig. 3A (left) shows the average eigenvalues across all scans, indicating that between 0.02 − 0.18*Hz*, the top-10 eigenvalues were very close and steadily increasing over frequencies, likely indicative of a 1/f-like pattern. CSDs at frequencies below 0.02*Hz* and above 0.18*Hz* have a higher spread in eigenvalues, indicating that these are more easily split into components that deviate from broadband connectivity.

**Figure 3.**
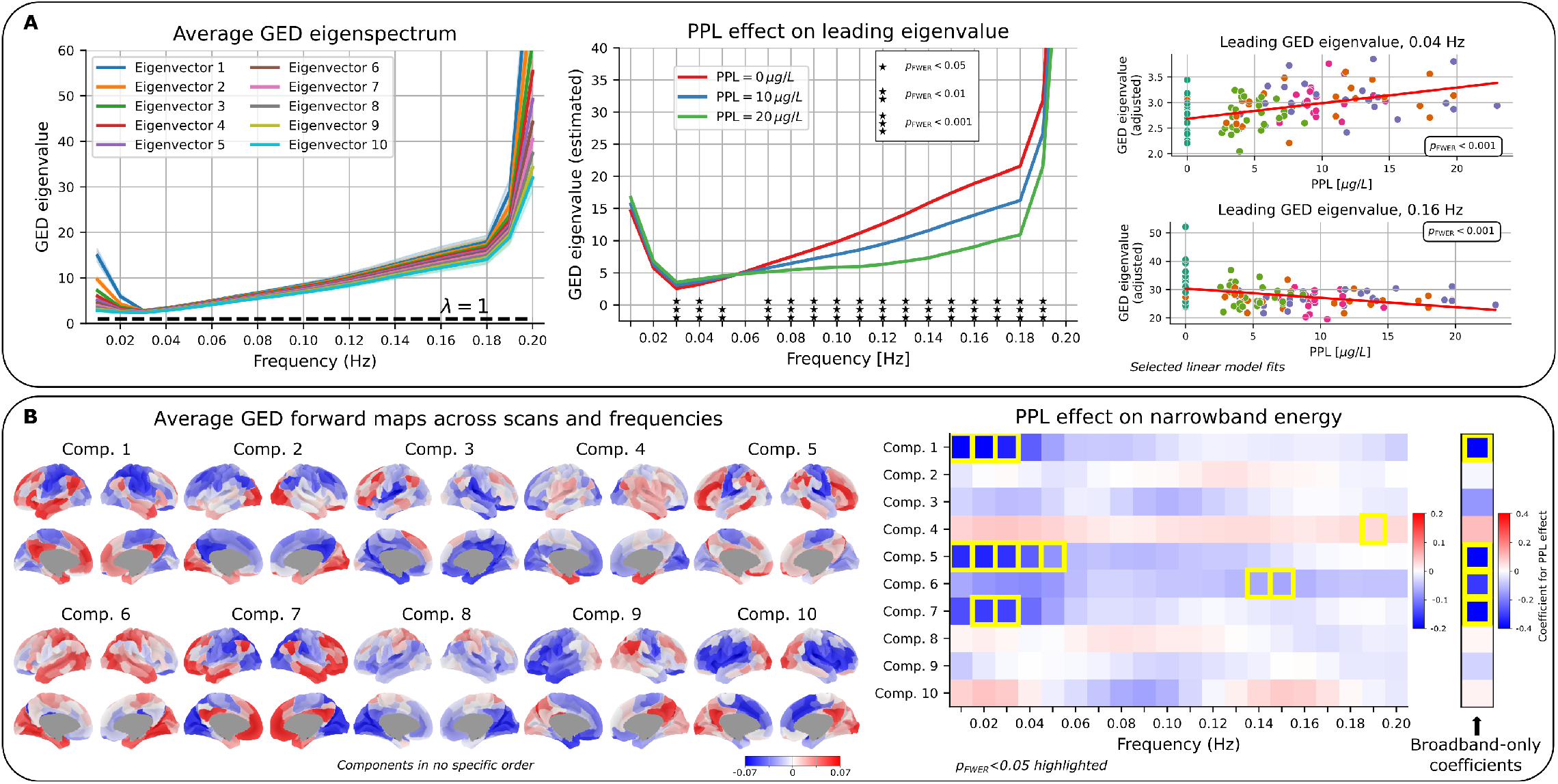
Generalized eigendecomposition (GED) on narrowband vs. broadband connectivity matrices and relation to psilocybin. **A**) Frequency-specific eigenvalues, which are interpretable as narrow-band/broadband variances, such that *λ >* 1 (horizontal line) indicates that the corresponding eigenvector explains more variance in narrowband than broadband connectivity. Associations between leading eigenvalues and PPL were assessed in a linear mixed effects model; estimates at selected PPL values are shown in the middle with significance indicated by asterisks. Selected linear model fits show log-power after regressing out nuisance variables and the subject-specific random effect (“adjusted”). **B**) Average GED forward maps were computed using dual-applied generalized Procrustes analysis (first across scans for each frequency, then across frequencies). The average maps were assessed for how much narrowband and broadband energy they explained in individual scans, and this coefficient was assessed for its relation to PPL. Statistically significant associations are highlighted with a yellow box.

We observed that PPL was significantly associated with the frequency-specific leading eigenvalue, although with a frequency-dependent sign (*f* = 0.03*Hz*: *r* = 0.52 [0.38, 0.64], *p*_FWER_ *<* 0.001; *f* = 0.15*Hz*: *r* = −0.42 [−0.56, −0.26], *p*_FWER_ *<* 0.001; Fig. 3A, middle). That is, PPL was negatively associated with the eigenvalue at lower frequencies (0.03 − 0.05*Hz*) and positively associated with the eigenvalue at higher frequencies (0.07 − 0.19*Hz*, most *p*_FWER_ *<* 0.001). This indicates that, at high PPL, narrowband connectivity becomes more distinct from broadband connectivity at lower frequencies, but converges to broadband connectivity at higher frequencies. An alternative explanation is that broadband connectivity is less influenced by lower frequencies following psilocybin administration, as also indicated by the decreased low-frequency univariate spectral power observed in Fig. 2A. We performed the same analysis on eigenvalues 2-10 and observed similar results (not shown), indicating that the shift in dominant frequency is not isolated to the leading eigencomponent.

To analyze the spatial representations computed by the GED, we matched scan- and frequency-specific forward maps using generalized Procrustes analysis (GPA) applied first across scans (within-frequency) and then across frequencies to obtain grand-average spatial components (Fig. 3B, left). Of note, since an eigenvector has arbitrary sign (i.e., it represents an axis), the forward map, representing the corresponding brain topography, may be sign-flipped without loss of information. Furthermore, the GPA does not account for differences in eigenvalue, and the ordering of components is therefore arbitrary. We observed average spatial patterns with high interhemispheric symmetry and typical unimodal/transmodal gradient patterns, suggesting that these are reflective of sensible underlying latent constructs.

We investigated how well the grand average spatial components explained energy in both broadband and narrowband scan- and frequency-specific connectivity by computing Rayleigh quotients and evaluating their relation to PPL (Fig. 3B, right). We observed that components 1, 5, 6, and 7 explained significantly less broadband connectivity as a function of PPL (comp. 1: *r* = −0.36 [−0.51, −0.20], *p*_FWER_ *<* 0.001). Components 1, 5, and 7 all segregated transmodal from unimodal cortices with only slightly varying loadings. These three components also had significantly reduced narrowband expression in the frequency range 0.01 − 0.05*Hz*, e.g., (comp. 1, *f* = 0.02*Hz*: *r* = −0.37 [−0.51, −0.20], *p*_FWER_ *<* 0.001). Meanwhile, component 6, which mainly segregated frontoparietal, temporal, and visual regions from the rest of the brain had significantly reduced narrowband expression at higher frequencies (*f* = 0.14*Hz*: *r* = −0.29 [−0.44, −0.12], *p*_FWER_ = 0.002). Most components expressed the same direction of change, except for visual-network dominated component 10, which indicated a statistically non-significant narrowband energy increase in low frequencies (∼ 0.02*Hz*) and a decrease at higher frequencies (∼ 0.11*Hz*), indicating a potential shift in the dominant frequency of the visual network.

## 4 Discussion

Here we present the first evaluation of the frequency-specific acute effects of psilocybin on BOLD fMRI activity and connectivity, examined in a cohort repeatedly scanned during the acute psilocybin experience. We found spatially widespread power decreases in low frequencies following psilocybin, strongest in frontoparietal/control, dorsal attention, and default mode networks, as well as evidence for psilocybin effects on frequency-specific connectivity, particularly related to the frontoparietal and visual networks. These results broaden our understanding of how psychedelics modulate BOLD signals and suggest that frequency-agnostic analyses of brain function/connectivity potentially mask frequency-specific effects of psilocybin.

Here we report the first characterization of frequency-specific psychedelic effects on BOLD fMRI. We found that spectral power decreased at slow frequencies (0.01 − 0.06*Hz*) following psilocybin administration, indicating a weakening of the BOLD response at these frequencies with a particular strong effect in transmodal networks including the control/frontoparietal, default, and dorsal attention networks. Our results suggest that psilocybin effects are likely confined to lower frequencies than the typical band of interest (0.01 − 0.08*Hz*), though future studies are needed to test this hypothesis. It is possible that this effect follows from a psilocybin-induced decrease in cerebral blood flow (CBF), which may in turn limit the hemodynamic response to neural activity (Carhart-Harris et al., 2012; Larsen et al., 2025; Lewis et al., 2017), potentially leading to a weakened BOLD signal variability. In a recent study by our lab (Larsen et al., 2025), CBF decreases following psilocybin were reported both globally and with strong effects in parietal, occipital, temporal, cingulate, and prefrontal cortices including evidence of psilocin-induced vasoconstriction of the internal carotid artery. The extent to which observed BOLD fluctuations are reflective of neuronal activity through the neurovascular coupling is a current topic of debate Epp et al., 2025, and it is possible that the reduced low-frequency BOLD signal variability induced by psilocin reflects vascular changes (static or dynamic) rather than decreased neuronal activity. Future studies should further investigate the neurovascular coupling mechanism and its perturbation by serotonergic drugs to disentangle neuronal from vascular activity.

We report an increase in spectral entropy, a metric that quantifies the flatness of the spectrum, initially supporting the theory of increased brain entropy (Carhart-Harris, 2018; Carhart-Harris et al., 2014; McCulloch et al., 2024). However, this finding appears to be driven by the observed reductions in low-frequency power (Fig. 2A). Reduced spectral power is directly related to a decrease in signal variance in these frequencies, which combined with a departure from the typical 1/f-like power distribution potentially indicates non-neural signal. Thus, an alternative explanation for entropy findings in psychedelic fMRI data, including temporal complexity or reduced network grouping of brain areas, is that these may reflect decreased low-frequency BOLD signal variability rather than increased neural “randomness”. A future study evaluating the relation between cerebral blood flow, frequency-specific BOLD signal power and connectivity patterns, and entropy quantifications could confirm this relation.

We provide the first report of narrowband connectivity patterns affected by psilocybin using the GED. We observed psilocybin effects on frequency-specific connectivity, particularly for the frontoparietal/control and visual networks, which have repeatedly been associated with psilocybin and other psychedelic drugs (La-Torraca-Vittori et al., 2025; Lord et al., 2019; Olsen et al., 2022). Our findings point towards a broadband reduction in connectivity along the transmodal/unimodal axis induced by psilocin. We believe that this result, when held in regard to the univariate control network power decrease at low frequencies (0.01 − 0.06*Hz*) primarily indicates a low-frequency reduction in transmodal connectivity. In contrast, the visual network (component 10) indicated a non-significant shift in its dominant frequency. These results reveal a complex frequency-specific connectivity shift exerted by psilocybin. Future studies should consider these results when determining filter bandwidths for, e.g., phase estimation which is predicated on narrowband data Olsen et al., 2024.

Head motion during BOLD scans can induce substantive confounding of functional connectivity estimates and is a particular challenge in psychedelic fMRI studies. Here, we sought to mitigate its effects through rigorous scan exclusion via motion criteria; denoising, including global signal regression (Ciric et al., 2017; Parkes et al., 2018); and inclusion of motion summary statistics as nuisance variables in statistical models. We expected increased motion to increase power over a wide range of frequencies due to their transient nature. Instead, our observation that PPL was negatively associated with low-frequency power indicates that these estimates are not driven by motion. Similarly, we assessed GED-based frequency-specific connectivity effects of psilocybin, which ensures that computed spatial patterns are maximally different from a reference covariance matrix. We assumed, as in previous studies (Rosso et al., 2025), that motion would be incorporated in the broadband covariance matrix. Combined with the indication that motion effects should be network-independent, we believe that we effectively mitigated the confounding effects of motion.

We suggested using the generalized Procrustes analysis to group and average forward maps, as computed via the GED, across subjects and frequencies. Eigenvectors are sign-invariant and the ordering of columns do not necessarily align across subjects and scans. Previous studies have either manually checked the consistency of information captured by the leading generalized eigenvectors and analyzed the most-often occurring pattern (Cohen, 2022), or disregarded the column ordering by exclusively evaluating the leading three generalized eigenvectors without correcting for column ordering (Rosso et al., 2025). As such, we believe the GPA is a principled way to combine generalized eigenvectors. Information about the relative importance of maps is lost in this procedure, but we showed that eigenvalues were generally very close (Fig. 3A). Furthermore, we retained and modeled only the leading 10 GED eigenvectors, although more had an eigenvalue above one, particularly at higher frequencies. Future studies could investigate methods for computing an average spatial map given variable input data dimensionality. In addition, future studies could compute GED eigenvectors and forward maps only at baseline or in a separate cohort. Here we wanted to ensure that potentially drug-altered spatial topographies would translate into the grand average.

We used linear mixed effects models to evaluate the association between PPL and spectral metrics, which assumed that this relation was without delay and that the effect of time was covered by PPL, i.e., upslope and downslope parts of the experiment corresponding to the same PPL were assumed to have the same outcome value. The variance in PPL across subjects differs throughout the experiment (Fig. 2A), which could indicate that a random slope between subject and drug would benefit our linear mixed effects models. However, we observed no non-linear interactions or temporal autocorrelation in residual plots. We also evaluated whether using unstructured covariance matrices in our modeling would improve type-1 error but found the opposite to be the case in synthetic experiments. Combined, we argue that the conventional linear mixed effects model is satisfactory in our case.

In conclusion, we found that psilocybin acutely reduces raw BOLD spectral power at slow frequencies, with particularly strong effects in transmodal networks. Furthermore, we observed reduced connectivity strength in unimodal/transmodal axis, particularly in low frequencies.

## Conflicts of interest

Brice Ozenne states that part of his salary while working on this project was covered by a grant from Novo Nordisk A/S, who did not have any involvement in the manuscript. MKM has received honoraria from Lobe Sciences as a scientific advisor. The other authors declare no competing interests.

## Author contributions

ASO, NuR, MG, and PMF conceived the study. ASO performed the experiments, constructed figures, and wrote the paper. KL, DE-WM, MKM, and PMF aided with analyses, interpretations, and visualization. BO provided statistical consultation. GMK, NuR, and PMF secured funding. All authors reviewed and approved the final manuscript.

## Data availability

All code pertaining to this study is made available on github: https://github.com/anders-s-olsen/Spectral-effects-of-psilocybin-fMRI. Data can be made available upon request and completion of a data sharing agreement.

## Notes

### Competing Interest Statement

The authors have declared no competing interest.

## References

Alamia, A., Timmermann, C., Nutt, D. J., VanRullen, R., & Carhart-Harris, R. L. (2020). DMT alters cortical travelling waves (V. van Wassenhove, T. E. Behrens, & D. M. Alexander, Eds.) [Publisher: eLife Sciences Publications, Ltd]. eLife, 9, e59784. 10.7554/eLife.59784

Barrett, F. S., Krimmel, S. R., Griffiths, R. R., Seminowicz, D. A., & Mathur, B. N. (2020). Psilocybin acutely alters the functional connectivity of the claustrum with brain networks that support perception, memory, and attention. NeuroImage, 218, 116980. 10.1016/j.neuroimage.2020.116980

Buzsáki, G., & Draguhn, A. (2004). Neuronal oscillations in cortical networks. Science (New York, N.Y.), 304 (5679), 1926–1929. 10.1126/science.1099745

Carhart-Harris, R. L. (2018). The entropic brain - revisited. Neuropharmacology, 142, 167–178. 10.1016/J.NEUROPHARM.2018.03.010

Carhart-Harris, R. L., Erritzoe, D., Williams, T., Stone, J. M., Reed, L. J., Colasanti, A., Tyacke, R. J., Leech, R., Malizia, A. L., Murphy, K., Hobden, P., Evans, J., Feilding, A., Wise, R. G., & Nutt, D. J. (2012). Neural correlates of the psychedelic state as determined by fMRI studies with psilocybin [Publisher: Proceedings of the National Academy of Sciences]. Proceedings of the National Academy of Sciences, 109 (6), 2138–2143. 10.1073/pnas.1119598109

Carhart-Harris, R. L., Leech, R., Hellyer, P. J., Shanahan, M., Feilding, A., Tagliazucchi, E., Chialvo, D. R., & Nutt, D. (2014). The entropic brain: A theory of conscious states informed by neuroimaging research with psychedelic drugs. Frontiers in Human Neuroscience, 8 (1 FEB), 20. 10.3389/FNHUM.2014.00020/BIBTEX

Ciric, R., Wolf, D. H., Power, J. D., Roalf, D. R., Baum, G. L., Ruparel, K., Shinohara, R. T., Elliott, M. A., Eickhoff, S. B., Davatzikos, C., Gur, R. C., Gur, R. E., Bassett, D. S., & Satterthwaite, T. D. (2017). Benchmarking of participant-level confound regression strategies for the control of motion artifact in studies of functional connectivity. NeuroImage, 154, 174–187. 10.1016/j.neuroimage.2017.03.020

Cohen, M. X. (2022). A tutorial on generalized eigendecomposition for denoising, contrast enhancement, and dimension reduction in multichannel electrophysiology. NeuroImage, 247, 118809. 10.1016/j.neuroimage.2021.118809

contributors, N. (n.d.). nilearn. 10.5281/zenodo.8397156

Delli Pizzi, S., Chiacchiaretta, P., Sestieri, C., Ferretti, A., Onofrj, M., Della Penna, S., Roseman, L., Timmermann, C., Nutt, D. J., Carhart-Harris, R. L., & Sensi, S. L. (2023). Spatial Correspondence of LSD-Induced Variations on Brain Functioning at Rest With Serotonin Receptor Expression. Biological Psychiatry: Cognitive Neuroscience and Neuroimaging, 8 (7), 768–776. 10.1016/j.bpsc.2023.03.009

Dryden, I. L., & Mardia, K. V. (2016, September). Statistical Shape Analysis, with Applications in R (1st ed.). Wiley. 10.1002/9781119072492

Epp, S. M., Castrillón, G., Yuan, B., Andrews-Hanna, J., Preibisch, C., & Riedl, V. (2025). BOLD signal changes can oppose oxygen metabolism across the human cortex [Publisher: Nature Publishing Group]. Nature Neuroscience, 1–12. 10.1038/s41593-025-02132-9

Faramarzi, A., Fooladi, M., Yousef Pour, M., Khodamoradi, E., Chehreh, A., Amiri, S., shavandi, M., & Sharini, H. (2024). Clinical utility of fMRI in evaluating of LSD effect on pain-related brain networks in healthy subjects. Heliyon, 10 (15), e34401. 10.1016/j.heliyon.2024.e34401

Godfrey, K., Luan, L. X., & Timmermann, C. (2025). Effects of psychedelics on human oscillatory brain activity. International Review of Neurobiology, 181, 171–202. 10.1016/bs.irn.2025.04.012

Golden, C. T., & Chadderton, P. (2022). Psilocybin reduces low frequency oscillatory power and neuronal phase-locking in the anterior cingulate cortex of awake rodents. Scientific Reports, 12 (1), 12702. 10.1038/s41598-022-16325-w

Gower, J. C. (1975). Generalized procrustes analysis. Psychometrika, 40 (1), 33–51. 10.1007/BF02291478

Haufe, S., Meinecke, F., Görgen, K., Dähne, S., Haynes, J.-D., Blankertz, B., & Bießmann, F. (2014). On the interpretation of weight vectors of linear models in multivariate neuroimaging. NeuroImage, 87, 96–110. 10.1016/j.neuroimage.2013.10.067

Ip, C.-T., Olbrich, S., de Bardeci, M., Monn, A., Ort, A., Smallridge, J. W., & Vollenweider, F. (2026). Psilocybin-induced alterations in EEG power, connectivity and network dynamics in healthy subjects: Correlations with subjective experience and implications for therapeutic applications. Progress in Neuro-Psychopharmacology and Biological Psychiatry, 145, 111626. 10.1016/j.pnpbp.2026.111626

Larsen, K., Lindberg, U., Ozenne, B., McCulloch, D. E., Armand, S., Madsen, M. K., Johansen, A., Stenbæk, D. S., Knudsen, G. M., & Fisher, P. M. (2025). Acute psilocybin and ketanserin effects on cerebral blood flow: 5-HT2AR neuromodulation in healthy humans. Journal of Cerebral Blood Flow and Metabolism: Official Journal of the International Society of Cerebral Blood Flow and Metabolism, 45 (7), 1385–1401. 10.1177/0271678X251323364

La-Torraca-Vittori, P., Tarchi, L., Arrigo, E., Lanterna, S., Tosi, E., Doose, A., Palesi, F., Pischedda, D., Ricca, V., Fusar-Poli, P., & Damiani, S. (2025). Knocking at the Doors of Perception: Relating LSD Effects on Low-Frequency Fluctuations and Regional Homogeneity to Receptor Densities in fMRIf. The European Journal of Neuroscience, 62 (10), e70338. 10.1111/ejn.70338

Lee, O. E., & Braun, T. M. (2012). Permutation Tests for Random Effects in Linear Mixed Models. Biometrics, 68 (2), 486–493. 10.1111/j.1541-0420.2011.01675.x

Lewis, C. R., Preller, K. H., Kraehenmann, R., Michels, L., Staempfli, P., & Vollenweider, F. X. (2017). Two dose investigation of the 5-HT-agonist psilocybin on relative and global cerebral blood flow. NeuroImage, 159, 70–78. 10.1016/j.neuroimage.2017.07.020

Lord, L. D., Expert, P., Atasoy, S., Roseman, L., Rapuano, K., Lambiotte, R., Nutt, D. J., Deco, G., Carhart-Harris, R. L., Kringelbach, M. L., & Cabral, J. (2019). Dynamical exploration of the repertoire of brain networks at rest is modulated by psilocybin. NeuroImage, 199 (April), 127–142. 10.1016/j.neuroimage.2019.05.060

Madsen, M. K., Fisher, P. M. D., Stenbæk, D. S., Kristiansen, S., Burmester, D., Lehel, S., Páleníček, T., Kuchař, M., Svarer, C., Ozenne, B., & Knudsen, G. M. (2020). A single psilocybin dose is associated with long-term increased mindfulness, preceded by a proportional change in neocortical 5-HT2A receptor binding. European Neuropsychopharmacology, 33, 71–80. 10.1016/j.euroneuro.2020.02.001

Madsen, M. K., Stenbæk, D. S., Arvidsson, A., Armand, S., Marstrand-Joergensen, M. R., Johansen, S. S., Linnet, K., Ozenne, B., Knudsen, G. M., & Fisher, P. M. (2021). Psilocybin-induced changes in brain network integrity and segregation correlate with plasma psilocin level and psychedelic experience. European Neuropsychopharmacology, 50, 121–132. 10.1016/j.euroneuro.2021.06.001

Markiewicz, C. J., Esteban, O., Goncalves, M., Poldrack, R. A., & Gorgolewski, K. J. (2026, March). fMRIPrep: A robust preprocessing pipeline for functional MRI. 10.5281/zenodo.18941091

McCulloch, D. E.-W., Knudsen, G. M., Barrett, F. S., Doss, M. K., Carhart-Harris, R. L., Rosas, F. E., Deco, G., Kringelbach, M. L., Preller, K. H., Ramaekers, J. G., Mason, N. L., Müller, F., & Fisher, P. M. (2022). Psychedelic resting-state neuroimaging: A review and perspective on balancing replication and novel analyses. Neuroscience & Biobehavioral Reviews, 138 (May), 104689. 10.1016/j.neubiorev.2022.104689

McCulloch, D. E.-W., Olsen, A. S., Ozenne, B., Stenbæk, D. S., Armand, S., Madsen, M. K., Knudsen, G. M., & Fisher, P. M. (2024, May). Navigating the chaos of psychedelic fMRI brain-entropy via multi-metric evaluations of acute psilocybin effects [ISSN: 2329-2164 Pages: 2023.07.03.23292164]. 10.1101/2023.07.03.23292164

Mitra, P. P., & Pesaran, B. (1999). Analysis of Dynamic Brain Imaging Data. Biophysical Journal, 76 (2), 691–708. 10.1016/S0006-3495(99)77236-X

Murray, C. H., Tare, I., Perry, C. M., Malina, M., Lee, R., & de Wit, H. (2022). Low doses of LSD reduce broadband oscillatory power and modulate event-related potentials in healthy adults. Psychopharmacology, 239 (6), 1735–1747. 10.1007/s00213-021-05991-9

Muthukumaraswamy, S. D., Carhart-Harris, R. L., Moran, R. J., Brookes, M. J., Williams, T. M., Errtizoe, D., Sessa, B., Papadopoulos, A., Bolstridge, M., Singh, K. D., Feilding, A., Friston, K. J., & Nutt, D. J. (2013). Broadband cortical desynchronization underlies the human psychedelic state [ISBN: 1517115183]. Journal of Neuroscience, 33 (38), 15171–15183. 10.1523/JNEUROSCI.2063-13.2013

Olsen, A. S., Brammer, A., Fisher, P. M., & Moerup, M. (2024, November). Uncovering dynamic human brain phase coherence networks [Pages: 2024.11.15.623830 Section: New Results]. 10.1101/2024.11.15.623830

Olsen, A. S., Lykkebo-Valløe, A., Ozenne, B., Madsen, M. K., Stenbæk, D. S., Armand, S., Mørup, M., Ganz, M., Knudsen, G. M., & Fisher, P. M. (2022). Psilocybin modulation of time-varying functional connectivity is associated with plasma psilocin and subjective effects. NeuroImage, 264, 119716. 10.1016/j.neuroimage.2022.119716

Parkes, L., Fulcher, B., Yücel, M., & Fornito, A. (2018). An evaluation of the efficacy, reliability, and sensitivity of motion correction strategies for resting-state functional MRI. NeuroImage, 171, 415–436. 10.1016/j.neuroimage.2017.12.073

Parra, L., & Sajda, P. (2003). Blind source separation via generalized eigenvalue decomposition. J. Mach. Learn. Res., 4 (null), 1261–1269. Retrieved January 30, 2026, from 10.5555/945365.964305

Pinheiro, J., & Bates, D. M. (2000). Mixed-Effects Models in S and S-PLUS. Springer-Verlag. 10.1007/b98882

Power, J. D., Barnes, K. A., Snyder, A. Z., Schlaggar, B. L., & Petersen, S. E. (2012). Spurious but systematic correlations in functional connectivity MRI networks arise from subject motion. NeuroImage, 59 (3), 2142–2154. 10.1016/j.neuroimage.2011.10.018

Power, J. D., Plitt, M., Laumann, T. O., & Martin, A. (2017). Sources and implications of whole-brain fMRI signals in humans. NeuroImage, 146, 609–625. 10.1016/j.neuroimage.2016.09.038

R Core Team. (2023). R: A language and environment for statistical computing. R Foundation for Statistical Computing. Vienna, Austria. https://www.R-project.org/

Rokem, A., Trumpis, M., & Pérez, F. (2009). Nitime: Time-series analysis for neuroimaging data, 68–75. 10.25080/WXDN0820

Roseman, L., Nutt, D. J., & Carhart-Harris, R. L. (2017). Quality of Acute Psychedelic Experience Predicts Therapeutic Efficacy of Psilocybin for Treatment-Resistant Depression. Frontiers in Pharmacology, 8, 974. 10.3389/fphar.2017.00974

Rosso, M., Fernández-Rubio, G., Keller, P. E., Brattico, E., Vuust, P., Kringelbach, M. L., & Bonetti, L. (2025). FREQ-NESS Reveals the Dynamic Reconfiguration of Frequency-Resolved Brain Networks During Auditory Stimulation [eprint: https://advanced.onlinelibrary.wiley.com/doi/pdf/10.1002/advs.20241 Advanced Science, 12 (20), p2413195. 10.1002/advs.202413195

Satterthwaite, T. D., Wolf, D. H., Ruparel, K., Erus, G., Elliott, M. A., Eickhoff, S. B., Gennatas, E. D., Jackson, C., Prabhakaran, K., Smith, A., Hakonarson, H., Verma, R., Davatzikos, C., Gur, R. E., & Gur, R. C. (2013). Heterogeneous Impact of Motion on Fundamental Patterns of Developmental Changes in Functional Connectivity During Youth. NeuroImage, 83, 10.1016/j.neuroimage.2013.06.045. https://doi.org/10.1016/j.neuroimage.2013.06.045

Schaefer, A., Kong, R., Gordon, E. M., Laumann, T. O., Zuo, X.-N., Holmes, A. J., Eickhoff, S. B., & Yeo, B. T. T. (2018). Local-Global Parcellation of the Human Cerebral Cortex from Intrinsic Functional Connectivity MRI. Cerebral Cortex, 28 (9), 3095–3114. 10.1093/cercor/bhx179

Schenberg, E. E., Alexandre, J. F. M., Filev, R., Cravo, A. M., Sato, J. R., Muthukumaraswamy, S. D., Yonamine, M., Waguespack, M., Lomnicka, I., Barker, S. A., & da Silveira, D. X. (2015). Acute Biphasic Effects of Ayahuasca. PloS One, 10 (9), e0137202. 10.1371/journal.pone.0137202

Schönemann, P. H. (1966). A generalized solution of the orthogonal procrustes problem. Psychometrika, 31 (1), 1–10. 10.1007/BF02289451

Subramanian, S., Reneau, T. R., Perry, D., Chacko, R., Laumann, T. O., Flavin, K., Horan, C., Schweiger, J., Metcalf, N., Lenze, E. J., Snyder, A. Z., Dosenbach, N. U. F., Nicol, G., & Siegel, J. S. (2025). Psilocybin’s acute and persistent brain effects: A precision imaging drug trial [Publisher: Nature Publishing Group]. Scientific Data, 12 (1), 941. 10.1038/s41597-025-05189-0

Tagliazucchi, E., Carhart-Harris, R., Leech, R., Nutt, D., & Chialvo, D. R. (2014). Enhanced repertoire of brain dynamical states during the psychedelic experience. Human Brain Mapping, 35 (11), 5442–5456. 10.1002/HBM.22562

Tagliazucchi, E., Roseman, L., Kaelen, M., Orban, C., Muthukumaraswamy, S. D., Murphy, K., Laufs, H., Leech, R., McGonigle, J., Crossley, N., Bullmore, E., Williams, T., Bolstridge, M., Feilding, A., Nutt, D. J., & Carhart-Harris, R. (2016). Increased Global Functional Connectivity Correlates with LSD-Induced Ego Dissolution. Current biology: CB, 26 (8), 1043–1050. 10.1016/j.cub.2016.02.010

Thomson, D. (1982). Spectrum estimation and harmonic analysis. Proceedings of the IEEE, 70 (9), 1055–1096. 10.1109/PROC.1982.12433

Timmermann, C., Roseman, L., Haridas, S., Rosas, F. E., Luan, L., Kettner, H., Martell, J., Erritzoe, D., Tagliazucchi, E., Pallavicini, C., Girn, M., Alamia, A., Leech, R., Nutt, D. J., & Carhart-Harris, R. L. (2023). Human brain effects of DMT assessed via EEG-fMRI [Publisher: Proceedings of the National Academy of Sciences]. Proceedings of the National Academy of Sciences, 120 (13), e2218949120. 10.1073/pnas.2218949120

Van Dijk, K. R. A., Sabuncu, M. R., & Buckner, R. L. (2012). The influence of head motion on intrinsic functional connectivity MRI. NeuroImage, 59 (1), 431–438. 10.1016/j.neuroimage.2011.07.044

Wall, M. B., Lam, C., Ertl, N., Kaelen, M., Roseman, L., Nutt, D. J., & Carhart-Harris, R. L. (2023). Increased low-frequency brain responses to music after psilocybin therapy for depression. Journal of Affective Disorders, 333, 321–330. 10.1016/j.jad.2023.04.081

Wu, C. W., Gu, H., Lu, H., Stein, E. A., Chen, J.-H., & Yang, Y. (2008). Frequency Specificity of Functional Connectivity in Brain Networks. NeuroImage, 42 (3), 1047–1055. 10.1016/j.neuroimage.2008.05.035

Yeo, B. T. T., Krienen, F. M., Sepulcre, J., Sabuncu, M. R., Lashkari, D., Hollinshead, M., Roffman, J. L., Smoller, J. W., Zöllei, L., Polimeni, J. R., Fischl, B., Liu, H., & Buckner, R. L. (2011). The organization of the human cerebral cortex estimated by intrinsic functional connectivity. Journal of Neurophysiology, 106 (3), 1125–1165. 10.1152/jn.00338.2011

Zang, Y.-F., He, Y., Zhu, C.-Z., Cao, Q.-J., Sui, M.-Q., Liang, M., Tian, L.-X., Jiang, T.-Z., & Wang, Y.-F. (2007). Altered baseline brain activity in children with ADHD revealed by resting-state functional MRI. Brain & Development, 29 (2), 83–91. 10.1016/j.braindev.2006.07.002

